# Speed-controlled visual stimuli modulate fish collective dynamics

**DOI:** 10.64898/2025.12.05.692523

**Authors:** Raj Rajeshwar Malinda, Saeko Takizawa, Akiyuki Koyama, Takayuki Niizato, Hitoshi Habe, Hiroaki Kawashima

## Abstract

Fish navigation is modulated by visual stimulation and associated parameters, such as size, speed, and orientation, yet the regulatory mechanisms remain unclear. Here, we experimentally investigate how systematically controlled stimulus speed influences the temporal and spatial collective dynamics of *Hemigrammus bleheri*. Using a simple and customisable top-down projection setup, we displayed clockwise-rotating black-dot patterns at three distinct speeds to simulate complex environmental motion cues. We quantified individual temporal responsiveness through angular movements, and evaluated group-level spatial organisation. Fish angular movements positively tracked the rotation direction of the visual patterns. Lower stimulus speeds elicited predominantly exploratory behaviour, while higher speeds induced a transition to strong motion-tracking behaviour, aligning fish orientation with the rotating patterns. Increasing stimulus speed also corresponded with greater nearest-neighbour distance, indicating reduced spatial cohesion. Notably, we observed transient schooling episodes not aligned with the rotational stimulus, reflecting periods in which internal social interactions outweighed stimulus-driven coordination. Using a newly defined free-schooling index, we show that such internally driven episodes were frequent at lower stimulus speeds but markedly reduced at higher speeds. Overall, these results demonstrate that speed-controlled visual stimuli modulate both individual navigation and group coherence, offering new insights into how visual cues shape fish collective behaviour dynamics.

## INTRODUCTION

Collective animal behaviour is a complex, multiscale phenomenon [1,2] that plays essential roles in biological systems [3–5]. These group-level patterns emerge from local interactions among individuals [1,2,6], and have been conceptualised through various modelling frameworks, including flocking in birds [7–9], swarming in insects [10,11], and schooling patterns in fish [12–16]. In aquatic animals, fish exhibit collective emergence through schooling and shoaling patterns that are amenable to quantitative observation and analysis [1,17–21]. These coordinated group-level responses support ecologically relevant functions including the ability to respond to prey-predator interactions [22,23], mating opportunities [24], foraging efficiency [25], swimming performance [26,27], and ultimately survival in natural habitats. Importantly, such behavioural phenotypes are modulated by external sensory cues, particularly vision and lateral-line mechanoreception (hydrodynamics sensing) [28–33], and are thought to widely influence on spatiotemporal coordination within the school [13,28,34–36].

Vision plays a vital and central role in coordinating [37] fish movement through motion and positional information that underpins visuo-motor coupling [38], alignment [26] and cohesion [39]. Visual perception enables individuals to estimate the heading, speed, and orientation of conspecifics, thereby regulating inter-individual spacing and temporal coordination [27,40–42]. Visual cues-mediated characteristics and parameters are crucial in navigational and coordinated animal behaviour, and drive various regulatory mechanisms in collective dynamics in fish [19,42–47]. Previous studies have explored both static visual cues [19], and dynamic illuminance-dependent effects [39,48,49], and have demonstrated that speed-dependent influence can alter directional movement in zebrafish [42,50,51].

Experiments employing robotic fish further suggest that the stimuli speed can modulate group coherence without substantially altering mean group speed, indicating that stimulus speed may function as a regulatory parameter in collective coordination [41]. Moreover, studies using virtual conspecifics have shown that decision accuracy of fish groups in zebrafish [50], preference behaviour to biological motions [51,52], different orientational behaviour [42], were related to the speed of visual cues. This study further argues that group size was found to impact decision accuracy, indicating a speed-accuracy trade-off mechanism in social information processing [50]. Collectively, these studies indicate that stimulus speed can diversify collective dynamics in fish movement [41,42,50–52]. However, a controlled, parametric examination of how speed-shaped visual cues modulate navigational behaviour, and how internal interactions may override stimulus-driven coordination, has not been fully explored.

From a methodological perspective, studies investigating visually mediated collective behaviour have employed a range of stimulus delivery methods, including side-view screens [19] and bottom-up projection systems [42,50,53]. Side-view methods are straightforward to implement, but they constrain the spatial geometry of visual cues, preventing uniform stimulus exposure across individuals and biasing the natural direction of movement. In contrast, bottom-up projection delivers visual cues directly onto the horizontal swimming plane, allowing more uniform stimulus exposure while preserving natural swimming freedom, and has therefore been used in immersive behavioural assays, including neurophysiological studies. However, the need for custom projector-tank assemblies limits the scalability of bottom-up systems, particularly in large-tank or aquaculture environments.

An alternative approach that retains the advantages of uniform stimulation across the horizontal swimming plane while avoiding the scalability constraints of tank-integrated projection assemblies is top-down projection. To date, however, applications of top-down projection in schooling fish studies have focused primarily on manipulating illuminance intensity rather than dynamic stimulus properties [39,48,49]. While these studies demonstrate that light levels modulate polarization, alignment, and cohesion in rummy-nose tetra species [39,48], recent large-tank implementations, including a pattern-based dynamic stimulus experiments with *Plecoglossus altivelis* in a 3 m × 3 m facility [44] show that dynamic visual cues can also be delivered at scale while preserving natural group movement. This indicates that top-down projection is not only scalable, but also suitable for analysing collective behaviour under ecologically relevant free-swimming conditions. However, how dynamic visual cues, particularly stimulus speed, shape navigational coordination and interact with internally maintained social interactions remains insufficiently clarified.

In this study, we first developed and validated a top-down projection system capable of delivering parametrically controlled dynamic visual stimuli across the swimming plane, providing a reproducible and practical framework for behavioural assays in freely schooling fish. Using this system, we systematically examined how variations in stimulus speed modulate temporal and spatial patterns of schooling behaviour in *H. bleheri*. To distinguish stimulus-driven coordination from internally maintained social interactions, we introduce a behavioural index that quantifies the relative contribution of internal versus visually induced alignment. This framework enables us to evaluate how external visual cues and intrinsic social dynamics jointly shape collective navigation, and introduces a top-down projection approach suitable for studying visually mediated social behaviour in freely swimming fish.

## RESULTS

### Validation of top-down projection system

We validated the top-down projection system for delivering dynamic visual stimuli to freely schooling fish (**Figure 1a**). This configuration enables visual stimuli to be projected onto the swimming plane while fish are recorded from above, allowing clear separation between the stimulus and body silhouettes using visible light only, without requiring infrared illumination or spectral filtering. The visual stimulus consisted of a clockwise rotating random-dot pattern, illustrated schematically in **Figure 1b**. The stimulus was controlled remotely, ensuring that no experimenter presence or disturbance influenced the fish during the behavioural assays. The visual stimulus was presented at three rotational speeds corresponding to slow (S1), medium (S2), and fast (S3) conditions, at approximately 17.2°, 34.3°, and 68.6° per second, respectively, which were selected to span the natural locomotion speed range of freely swimming *Hemigrammus bleheri*. The stimulus presentation and behavioural recording were conducted at 60 frames per second (fps). Each stimulus condition, including the control, was presented three times per day across three consecutive days, with the central 35-second segment from each 40-second trial used for quantitative analyses. The detailed experimental protocol, including stimulus presentation timing and recording phases, is provided in **Methods and Supplementary Figure 1**.

**Figure 1:**
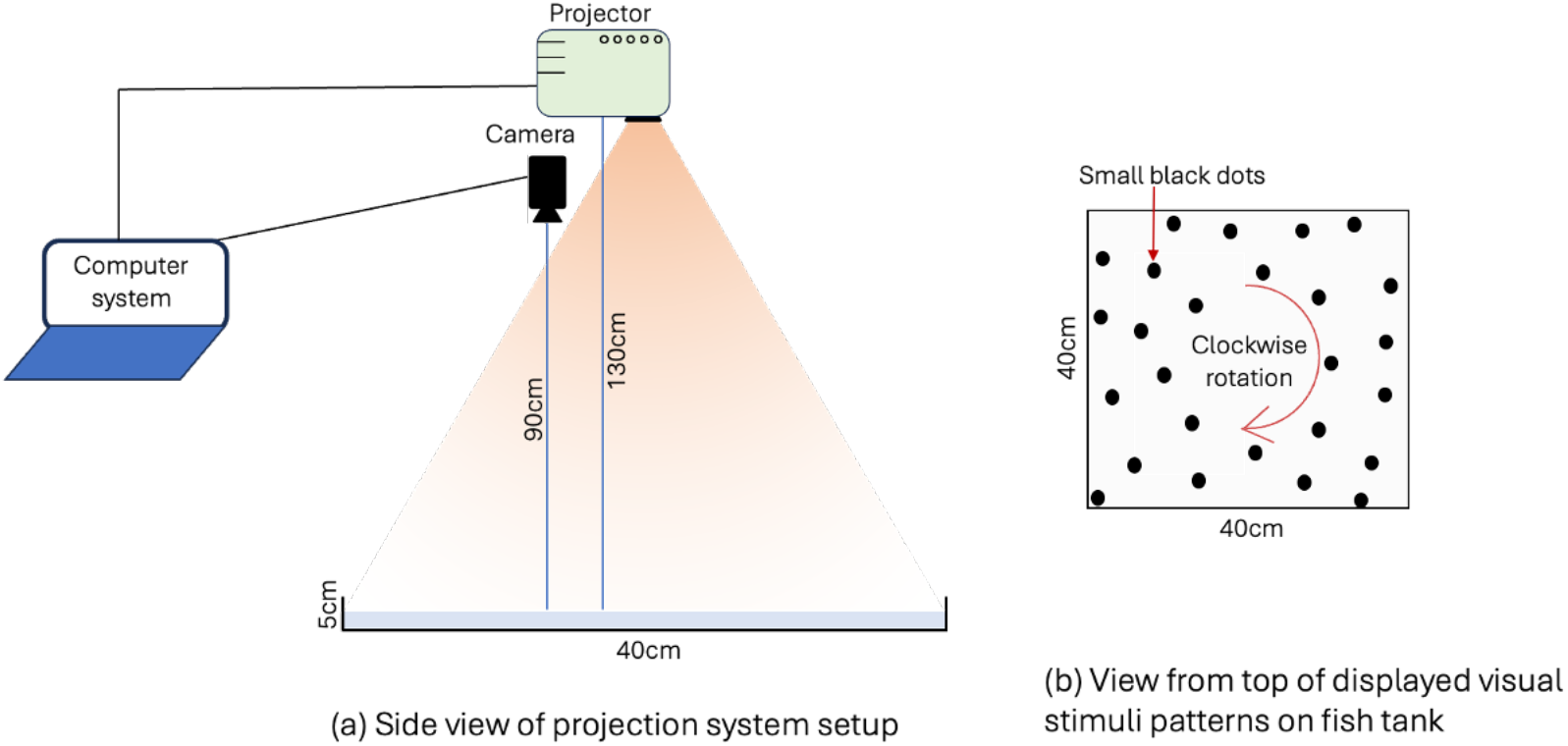
Top-down projection system and schematic representation of the visual stimulus. **(a)** Side-view of the experimental setup, including the projector, overhead camera, and control computer used to deliver visual stimuli to the fish tank. **(b)** Schematic illustration of the visual stimulus pattern displayed on the swimming plane, showing the spatial layout and clockwise rotational motion of the pattern.

Individual fish and the projected random-dot stimulus were detected using two separate YOLO11-based models [54,55]. Fish were detected in each frame and continuously tracked to maintain individual identities across frames, with manual corrections required in less than 1% of frames. The bounding-box centres were used to generate two-dimensional trajectories. The rotation centre of the projected dot pattern (hereafter referred to as the centre of rotation, COR) was estimated from the detected dot positions in each frame; because this stimulus rotated as a rigid pattern, its centre position **p**_*c*_ remained constant within each video. The resulting dataset provided time series of individual fish positions **p**_*i*_(*t*), from which instantaneous velocities were obtained after temporal smoothing with a 9-frame (≈ 0.15 s) moving average, and relative orientations to the COR were computed for subsequent analyses.

Based on these trajectories, we calculated the mean angular velocity (*ω*_*i*_, °/s) of each individual *i* relative to the COR to quantify stimulus-following behaviour (**Supplementary Figure 2**). In each trial, *ω*_*i*_ was defined as the time-averaged angular velocity for each individual (see *Methods*). Positive *ω*_*i*_ values indicated clockwise motion, corresponding to alignment with the rotating stimulus. The distributions of *ω*_*i*_ across the three speed conditions (S1−S3) showed a clear upward shift with increasing stimulus speed (**Figure 2**). Median *ω*_*i*_ values were 9.0°, 15.3°, and 33.4°/s for S1, S2, and S3, respectively, indicating that the fish’s angular responses scaled with the projected stimulus speed. Kruskal−Wallis test confirmed significant differences among conditions (*H* = 104.2, *p* = 2.0 × 10^-22^), and post-hoc Dunn’s tests with Bonferroni correction showed that all speed conditions differed significantly from control (S1: *p* = 2.6 × 10^-3^; S2: *p* = 4.3 × 10^-7^; S3: *p* = 8.6 × 10^-23^). These results demonstrate not only that stimulus speed strongly modulated individual angular velocity, but also that the projection system effectively elicited and captured speed-dependent visual responses in freely swimming fish.

**Figure 2:**
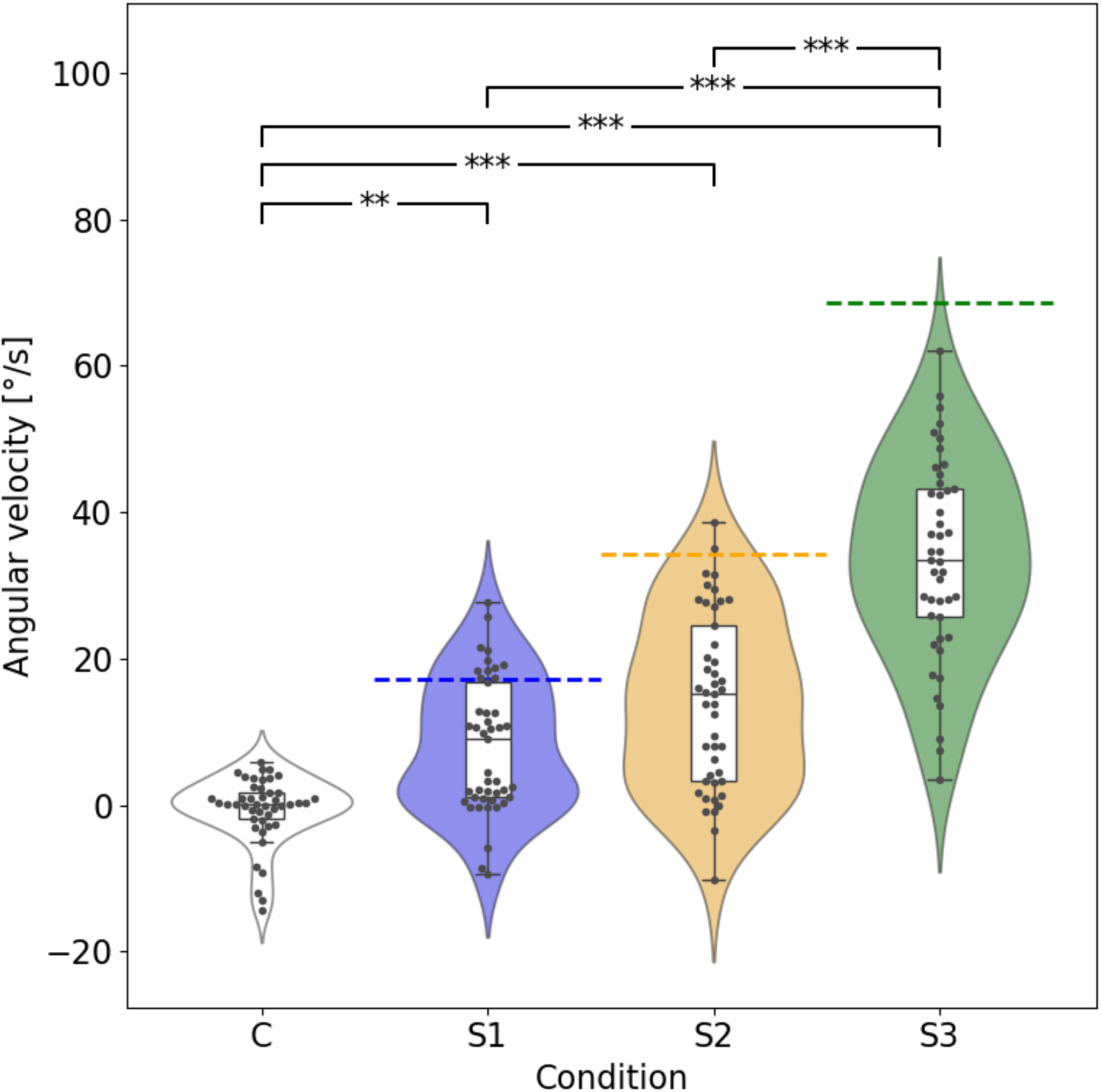
Angular velocity of individual fish under different stimulus speeds. Violin plots illustrate the distribution of individual angular velocities (°/s), and box plots indicate the median and interquartile ranges. Dashed lines represent the projected angular velocities (°/s) of the rotating visual patterns for each condition. Asterisks denote statistically significant pairwise differences among conditions based on Dunn’s post-hoc tests with Bonferroni correction (* p < 0.05, ** p < 0.01, *** p < 0.001).

To further examine the behavioural dynamics induced by speed-dependent visual stimuli, we next conducted a detailed temporal and spatial analysis to evaluate the validity of the stimulus speeds. We also investigated how stimulus following and free schooling alternated over time by introducing a new behavioural index, the free-schooling index (FSI).

### Temporal and spatial analyses of schooling behaviour

To evaluate how stimulus speed shaped collective coordination, we first examined the distribution of individual angular velocities. As shown in **Figure 2**, the distributions exhibited clear bimodality under the slower stimulus conditions, indicating that both responding and non-responding individuals were present within the group during these trials. Detailed inspection of individual trajectories revealed that fish alternated dynamically between stimulus-following and non-following phases within a single trial. In particular, trajectories in the fastest condition (S3) approached a nearly circular path, whereas those in S1 and S2 showed transitional patterns between exploratory and rotational motion (representative examples are provided in **Supplementary Figure 3**).

To isolate periods of active following, we filtered frames in which the swimming direction was within ±45° of the tangential direction relative to the centre of rotation (COR), corresponding to motion aligned with the rotating stimulus (see also **Supplementary Figure 2**). Under this filtering condition, the distributions of mean angular velocities became more unimodal and shifted toward higher values, especially in S1 and S2 (**Figure 3**). When comparing the medians of filtered mean angular velocities (20.1°, 29.0°, and 42.4°/s for S1, S2, and S3, respectively), the median for S1 slightly exceeded the stimulus speed, whereas that for S2 was marginally lower, and that for S3 was markedly lower than the fastest stimulus. These results indicate that the natural range of visually guided locomotion in *H. bleheri* lies between the S1 and S2 regimes, whereas the S3 condition exceeded that range.

**Figure 3:**
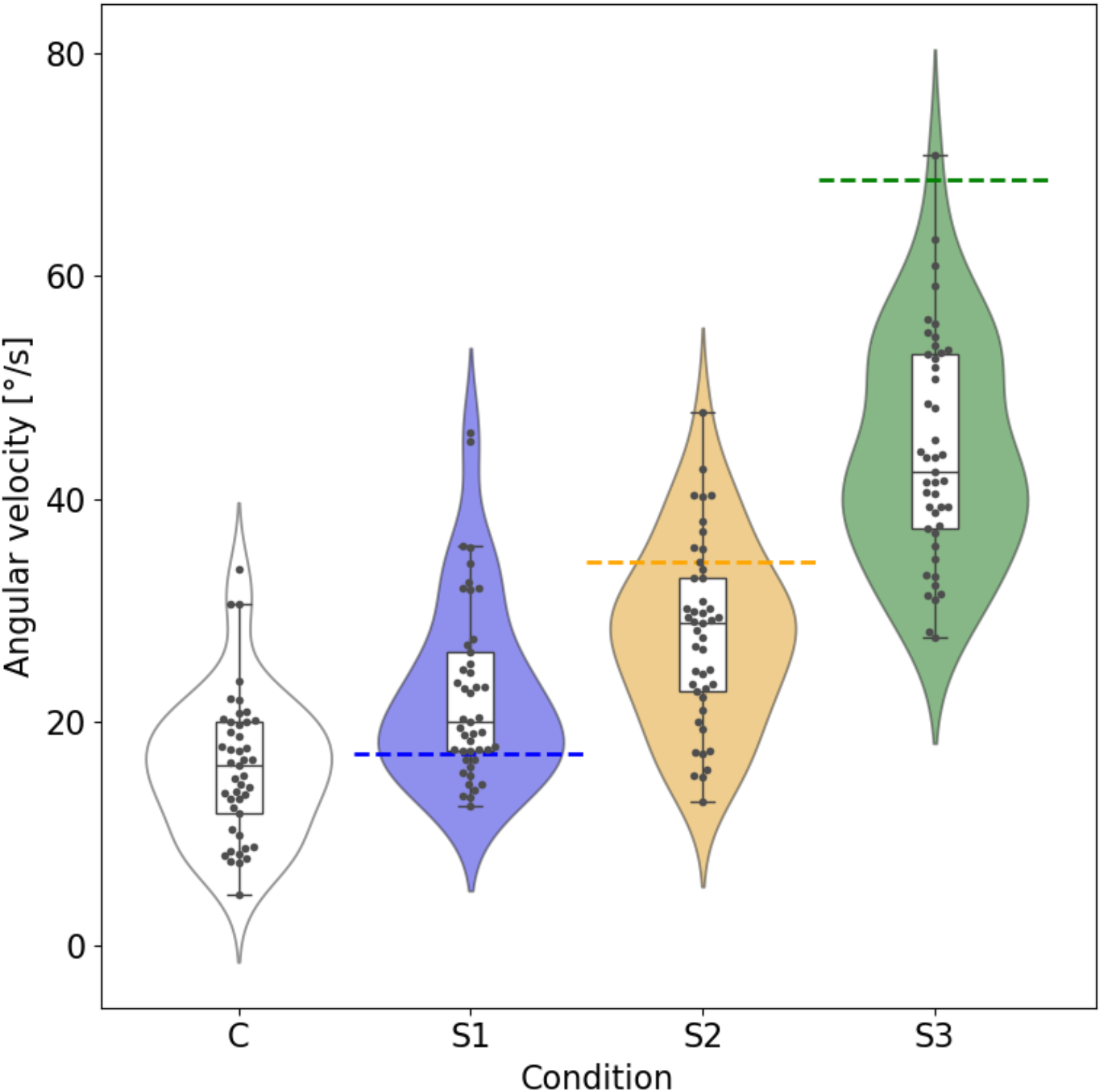
Angular velocity of individual fish during stimulus-aligned rotational periods. Violin plots illustrate the distribution of individual angular velocities (°/s) after filtering frames to include only motion within ±45° of the clockwise tangential direction around the COR, and box plots indicate the median and interquartile ranges. Dashed lines represent the projected angular velocities (°/s) of the rotating visual patterns for each stimulus condition.

We next analysed spatial organisation to determine how speed-dependent stimuli influenced group structure. Under control conditions, fish displayed exploratory movement near the tank boundaries, whereas under rotating stimuli, their trajectories became more circular and coherent around the COR (**Supplementary Figure 3**). The nearest-neighbour distance (NND) also changed systematically with stimulus speed (**Figure 4**). A Kruskal– Wallis test revealed significant differences among conditions (*H* = 46.7, *p* = 4.0 × 10^-10^). Post-hoc comparisons showed that NND increased significantly in S1 (*p* = 2.1 × 10^-2^), S2 (*p* = 4.1 × 10^-5^), and S3 (*p* = 2.1 × 10^-10^) compared with control, and S3 showed significantly larger NND than S1 (*p* = 1.3 × 10^-3^). The enlarged NND at higher stimulus speeds suggests a reduction in local alignment and cohesion, reflecting a shift toward stimulus-driven rather than socially maintained organization. Together, these results demonstrate that speed-controlled visual stimuli modulated both temporal and spatial aspects of schooling, delineating a behavioural transition from exploratory to stimulus-following motion.

**Figure 4:**
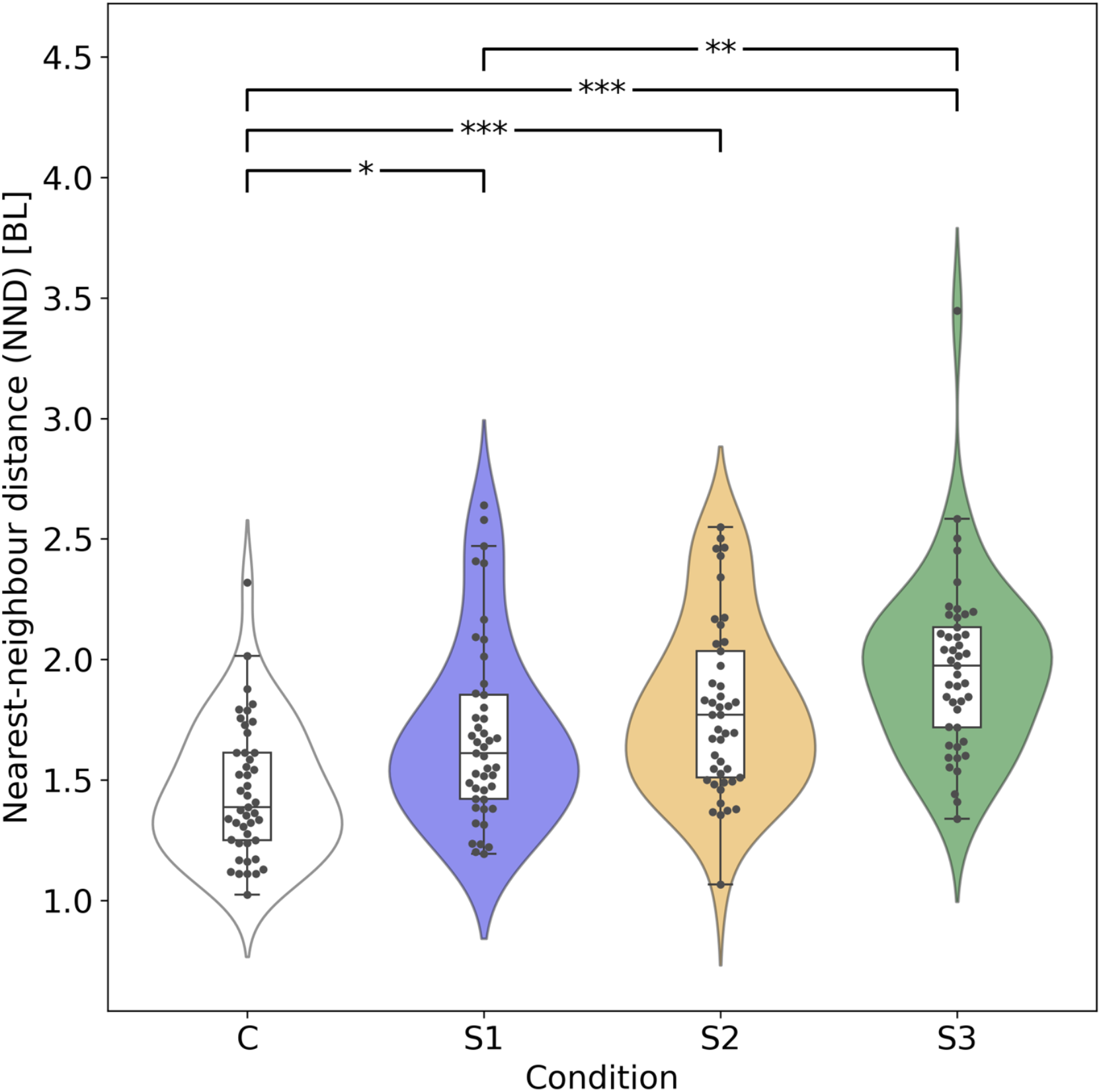
Nearest-neighbour distance under different stimulus speeds. Violin plots illustrate the distribution of nearest-neighbour distances expressed in body lengths (BL), and box plots indicate the median and interquartile ranges. Nearest-neighbour distance increased systematically with stimulus speed. Asterisks denote statistically significant pairwise differences among conditions based on Dunn’s post-hoc tests with Bonferroni correction (* p < 0.05, ** p < 0.01, *** p < 0.001).

### Quantifying internal–stimulus balance in schooling behaviour

To quantify the balance between visually driven rotational behaviour and stimulus-independent schooling, we introduced a free-schooling index (FSI). Directional alignment is commonly measured through the velocity correlation between neighbouring individuals, typically expressed as a directional correlation (cosine similarity) of normalised velocity vectors. In the present study, because fish swam within a confined tank, we focused on the alignment with only each nearest neighbour and defined, for each individual *i*,

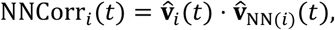

Where 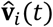 ) is the normalised velocity of fish *i*, and NN(*i*) denotes its nearest neighbour. Similarly, we quantified alignment with the visual stimulus by computing the directional correlation between the fish velocity and the tangential direction of the rotating pattern,

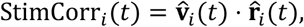

where 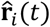 is the unit tangential direction of the stimulus evaluated at the position of individual *i* (see **Supplementary Figure 2**). In control trials, where no visual stimulus was presented, the same tangential direction field around the tank centre was used to compute StimCorr_*i*_(*t*), ensuring that the definition of FSI remained consistent across all conditions. The instantaneous free-schooling index was then defined as,

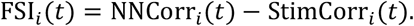

Positive values of FSI_*i*_(*t*) indicate periods when neighbour-based alignment exceeded stimulus-based alignment. Values near zero indicate that the two alignments were similar in magnitude, a situation that may arise when both alignments are strong or when both are weak.

Because our aim was to characterise each individual’s overall behavioural tendency within a trial, we computed a time-averaged index,

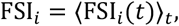

which is the value shown in **Figure 5**. Higher FSI_*i*_ values correspond to longer durations in which neighbour-based coordination dominated over stimulus-driven alignment, whereas lower values indicate trials in which stimulus-based alignment was comparable to or exceeded neighbour alignment.

**Figure 5:**
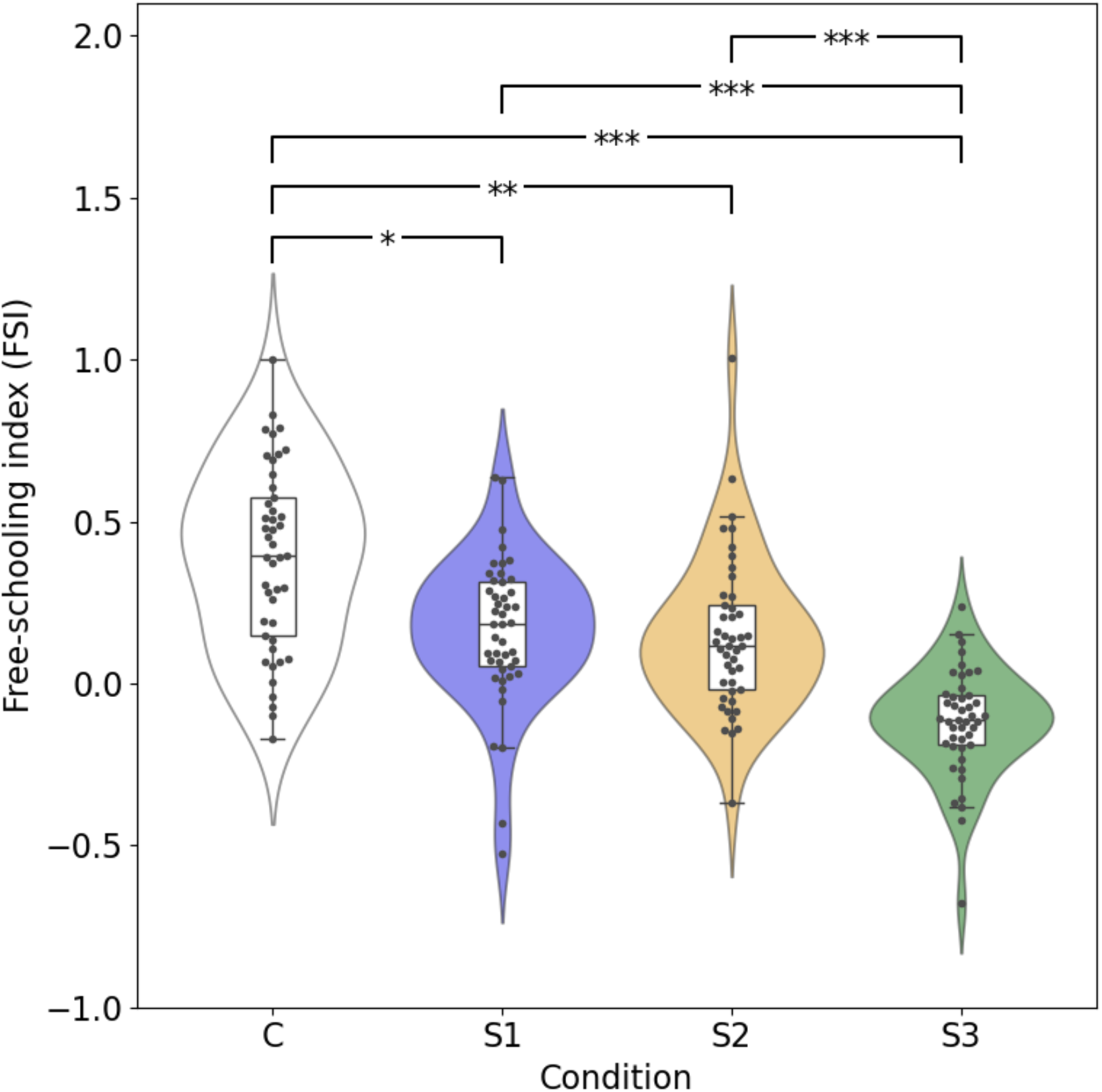
Free-schooling index (FSI) under different stimulus speeds. Violin plots illustrate the distribution of the free-schooling index (FSI), and box plots indicate the median and interquartile ranges. Higher FSI values reflect periods in which neighbour-based alignment exceeded stimulus-based alignment, reflecting internally maintained schooling behaviour. FSI decreased systematically with increasing stimulus speed, with the lowest values observed under the fastest condition (S3). Asterisks denote statistically significant pairwise differences among conditions based on Dunn’s post-hoc tests with Bonferroni correction (* p < 0.05, ** p < 0.01, *** p < 0.001).

As shown in **Figure 5**, FSI decreased systematically with increasing stimulus speed. Control trials exhibited the highest values, reflecting frequent free-schooling episodes, whereas S1 and S2 showed progressively reduced values. S3 produced the lowest FSI, indicating that the fastest stimulus strongly constrained the group to follow the imposed rotational motion. These differences were supported by statistical analysis: a Kruskal–Wallis test revealed a significant effect of stimulus speed on FSI (*H* = 73.8, *p* = 6.5 × 10^-16^), and Dunn’s post-hoc tests with Bonferroni correction showed that all stimulus conditions differed significantly from control (S1: *p* = 2.4 × 10^-2^; S2: *p* = 2.2 × 10^-3^; S3: *p* = 1.8 × 10^-16^). Together, these results demonstrate that increasing stimulus speed suppresses internally maintained schooling and promotes stronger stimulus-following behaviour.

## METHODS

### Fish maintenance and experimental setup

Rummy-nose tetra (*Hemigrammus bleheri*), with an average standard body length of approximately 3–5cm, were obtained from a commercial aquarium fish supplier. Fish were maintained under standard laboratory conditions in a 20 × 10 × 20 cm holding tank at 27 ± 1 °C, with the room temperature kept at 25 ± 1 °C. They were fed once daily in the morning at a fixed time using an automated feeder attached to the tank.

Fish were tested in a shallow square acrylic tank (40 × 40 × 5 cm) with a white matte interior surface, filled with 2–3 cm of water. A digital projector (EPSON EB-W06, WXGA, 3700 lm) was mounted approximately 130 cm above the tank to present visual stimuli onto the bottom surface. A video camera (ELP-USBFHD08s-MFV, 1080p, 60 fps) fitted with a fixed-focal-length lens (Kowa LM5JCM-WP, 5 mm) was positioned approximately 90 cm above the tank to record fish behaviour from above. Both devices were connected to a nearby laptop computer (CPU: Intel Core i9-13980HX; GPU: NVIDIA GeForce RTX 4080; RAM: 64GB DDR5; storage: 1TB SSD), which controlled stimulus presentation and video acquisition.

### Stimulus design and protocol

The visual stimuli consisted of small black dots scattered on a white background, rotating clockwise around a fixed centre of rotation (COR) at three discrete angular speeds (S1–S3). The COR was positioned approximately at the centre of the fish tank. Stimulus presentation followed a Latin-square protocol across conditions, as illustrated in **Supplementary Figure 1**, to minimize potential order or learning effects. Each stimulus period lasted 40 s, followed by a stimulus-off control period with a uniform white background, and the centre 35 s of each period was used for analysis.

### Experimental procedure

Experiments were conducted on three consecutive days at approximately the same time each morning. On each experimental day, five fish were randomly selected and transferred to the experimental tank approximately 35 ± 5 min before the experiment to allow acclimation.

After acclimation, three cycles were conducted per day. Each cycle began with a control period (C), followed by the three stimulus speeds (S1–S3) presented once each, with a blank (no-pattern) period inserted between successive stimulus presentations. Across the three cycles, the order of S1–S3 followed a Latin-square schedule, ensuring that each stimulus condition was presented exactly three times per day (see **Supplementary Figure 1**). A uniform white projection (without black dots) was used for both control and blank periods. Day 1 had an additional blank period after each cycle and 5-minute break with the projector off between cycles, while cycles continued without turning off the projector on days 2 and 3.

A total of 12 fish was involved in the three-day experiments, with some individuals participating on multiple days. No fish mortality occurred during or after the experiments. All sessions were recorded at 60 frames per second (fps) for subsequent analysis.

### Tracking analysis

All the recorded videos were processed to extract two-dimensional trajectories of individual fish. Detection was performed using two separately fine-tuned YOLO11 models [54]: one for fish and one for stimulus dots. Fish identities were maintained across frames using the BotSort tracking algorithm included in the YOLO11 framework [54,55]. Automatic tracking errors (e.g., ID swaps and missed detections) were manually corrected using a custom GUI tool.

Fish trajectories **p**_*i*_(*t*), defined as the centres of the tracked bounding boxes of individual *i*, were smoothed using a 9-frame (≈ 0.15 s) moving average. Velocities **v**_*i*_(*t*) were computed by framewise differentiation. The position of the centre of rotation (COR), **p**_*c*_, was estimated for each video from the detected stimulus dot patterns.

### Derivation of metrics and statistical analysis

Individual angular velocity and its time average were computed as,

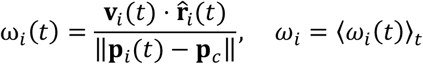

where 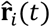 is the unit tangential direction of the stimulus evaluated at the position of individual *i*, obtained by rotating **p**_*i*_(*t*) − **p**_*c*_ by 90° clockwise and normalising it to unit length (see **Supplementary Figure 2**). To ensure stable computation, frames in which ‖**p**_*i*_(*t*) − **p**_*c*_‖ < 50 pixels were excluded from the analysis.

The average body length of fish in the recorded images was approximately 75-80 pixels; a value of 75 pixels was used as 1 body length (BL) when expressing nearest-neighbour distance (NND) in **Figure 4**.

For the computation of each individual’s NNCorr, StimCorr, and FSI, frames with ‖**v**_*i*_(*t*)‖ = 0 were treated as invalid and omitted from averaging. Differences among the four conditions (control, S1–S3) were assessed using the Kruskal–Wallis test, followed by Dunn’s post-hoc pairwise comparisons with Bonferroni correction. Significance thresholds were set at *p* < 0.05, *p* < 0.01, and *p* < 0.001 [56,57].

## DISCUSSION

Our study validated a top-down visual projection system (**Figure 1**) that provides an easy to set up, inexpensive, and non-invasive platform for analyses of collective behaviour in aquatic animals. This configuration enables visual stimuli to be projected directly onto the swimming plane while fish are recorded from above. Utilizing recent advances in multi-object detection and tracking [54,55], the present setup ensures clean separation between stimulus patterns and fish silhouettes with only minimal manual correction, eliminating the need for infrared illumination or spectral filtering [39,44]. The system supports customised and reproducible delivery of dynamic visual stimuli and is readily scalable from small laboratory arenas to large aquaculture facilities [58]. Owing to its modular structure, the platform can be combined with additional components when required, such as closed-loop intervention with real-time tracking [59], making it broadly applicable for studying visually mediated navigation across diverse experimental and ecological contexts.

Using this projection system, we characterised the temporal and spatial aspects of collective dynamics in *Hemigrammus bleheri* under systematically varied stimulus speeds. Controlled top-down projection induces clear, speed-dependent shifts in directional navigation, often overriding the intrinsic tendency of fish to explore near the tank boundary (**Supplementary Figure 3**) [46]. The distributions of angular velocity showed a systematic upward shift with stimulus speed (**Figure 2**), yet the medians of unfiltered responses did not match the projected rotation rates. This discrepancy can be explained that, the bimodal distributions in S1 and S2 indicate heterogeneous individual responsiveness, likely arising from individual differences in visual sensitivity, behavioural state, or moment-to-moment variation in attention[17,18,39,60].

Moreover, the angular-velocity analysis was restricted to frames in which individuals moved in the stimulus direction, S1 slightly exceeded the stimulus speed while S2 showed only a modest lag (**Figure 3**). These analyses indicate that heterogeneity, rather than insufficient stimulus strength [61], shaped the distributions at moderate speeds. However, S3 exhibited a pronounced lag even during responding frames, suggesting that the fastest rotation approached biomechanical or physiological limits of visually guided locomotion. Spatial structure also shifted with stimulus speed: nearest-neighbour distance increased systematically under faster rotation (**Figure 4**) [40,41], while group-level coherence was generally maintained. Together, these results demonstrate that visual motion cues can modulate both individual behaviour and group-level spacing in a markedly speed-dependent manner, while overall group cohesion remains largely preserved.

Notably, stimulus speed appeared to shift the balance between stimulus-following and neighbour-driven, stimulus-independent coordination. To characterise this balance, we introduced the free-schooling index (FSI) (**Figure 5**). Operationally, FSI is defined as the time-averaged difference between nearest-neighbour alignment and stimulus-based alignment, thereby capturing the extent to which individuals engaged in neighbour-driven rather than stimulus-driven coordination. This metric complements established measures such as polarisation by incorporating information about both local social interactions and alignment to an external visual cue [39,48,49,51]. In our data, FSI decreased systematically with increasing stimulus speed, with S3 showing the lowest values, revealing that fasters rotations reduced the prevalence of stimulus-independent schooling (**Figure 5**). These findings demonstrate that visual motion cues shape not only directional alignment but also the balance between internally maintained and stimulus-driven coordination [40,42,51].

In conclusion, the top-down projection-based system evaluated in this study provides a flexible, non-invasive, and scalable framework for experimentally probing visually mediated navigation in fish. Our results show that systematic variation in stimulus speed strongly modulates both individual and group-level behaviour, highlighting the role of dynamic visual motion cues as potent drivers of coordinated motion. The reproducibility and modularity of the system make it well suited for extension to closed-loop stimulation, comparative studies across species, and investigations into how different visual attributes, such as pattern geometry and light intensity, as well as their interactions with motion cues, shape collective dynamics. More broadly, this approach provides a versatile platform for dissecting the mechanisms of visually guided collective behaviour and offers a foundation for applications ranging from basic behavioural research to aquaculture and other applied settings.

## ETHICS

All experiments in this study were conducted in accordance with the guidelines of the departmental ethics committee of the Graduate School of Information Science, University of Hyogo, Japan, under the ethical approval number UHIS-EC-2024-004.

## DATA ACCESSIBILITY

All datasets and scripts will be available publicly upon the final publication.

## SUPPLEMENTARY DATA SHEET

**Supplementary Figure 1:**
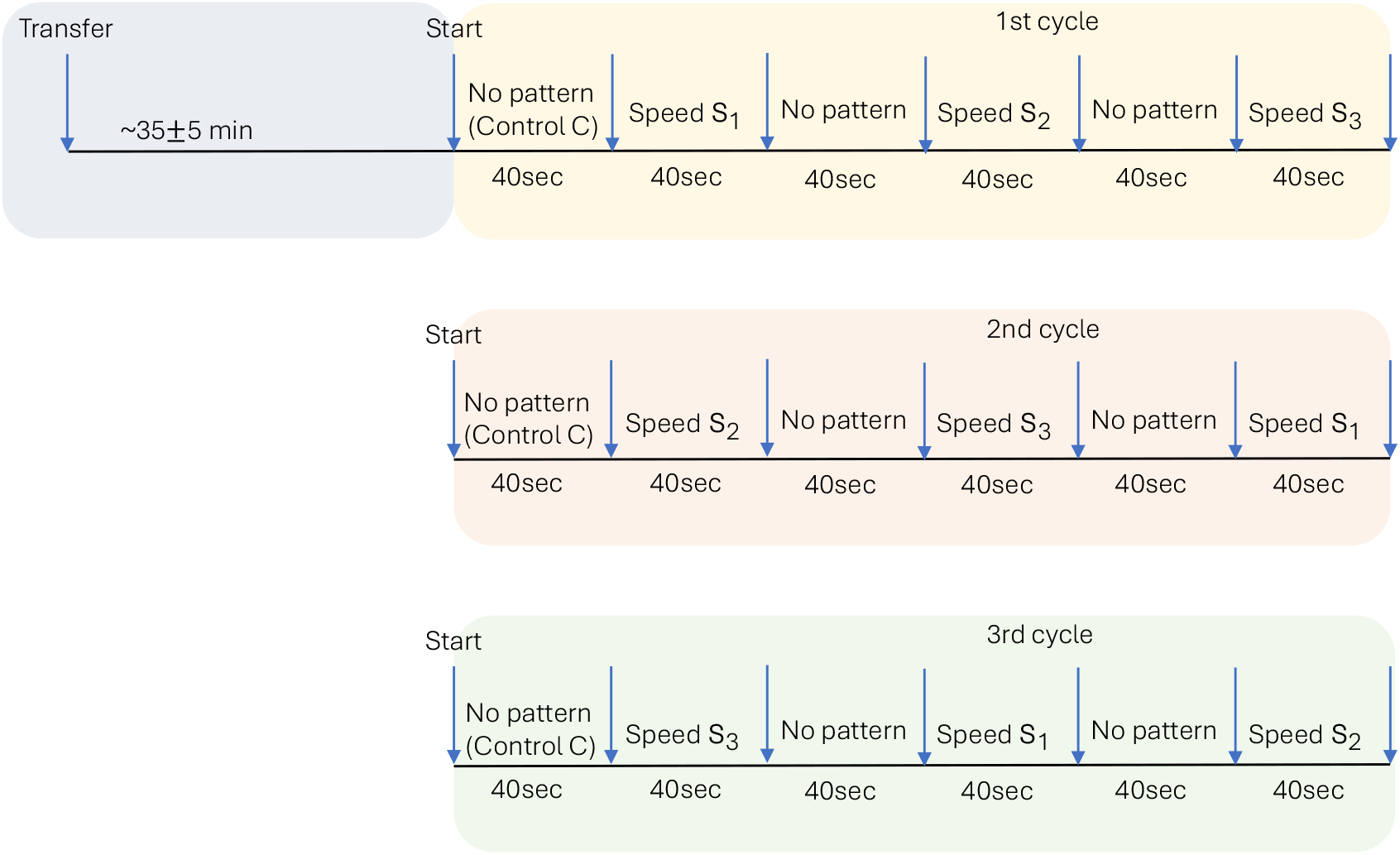
Experimental protocol used within each experimental day. The three stimulus speed conditions (S1, S2, and S3) were presented once per cycle in a Latin-square order to minimise sequence-dependent learning or habituation effects. Each stimulus period lasted 40 s and was separated by a blank (no-pattern) interval. A uniform white background was used during both control and blank phases. This three-cycle structure was repeated each day.

**Supplementary Figure 2:**
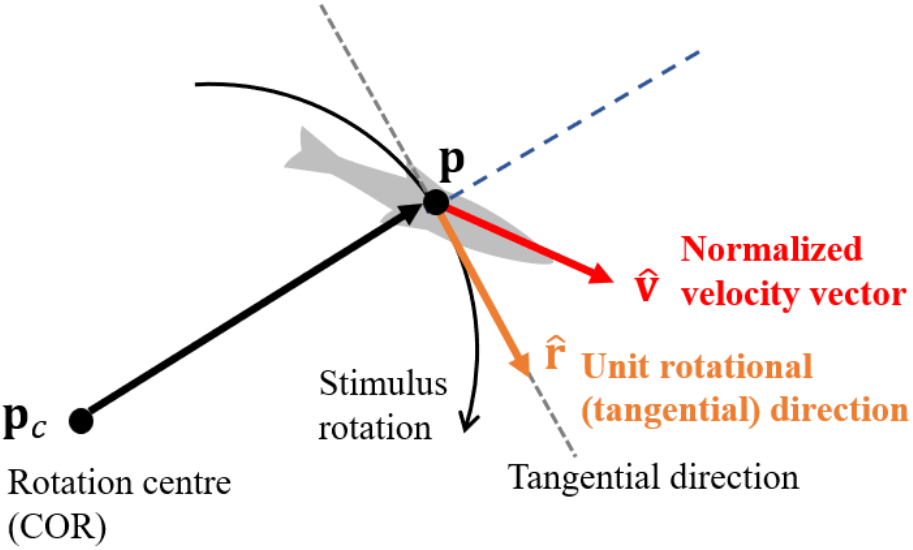
Illustration of the geometric quantities used in the analysis. **p**_c_ denotes the rotation centre (COR) of the projected visual stimulus, **p** represents the position of fish. 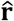 is the unit rotational (tangential) direction of the clockwise rotating stimulus evaluated at **p**, and 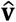 denotes the normalized velocity vector of the fish.

**Supplementary Figure 3:**
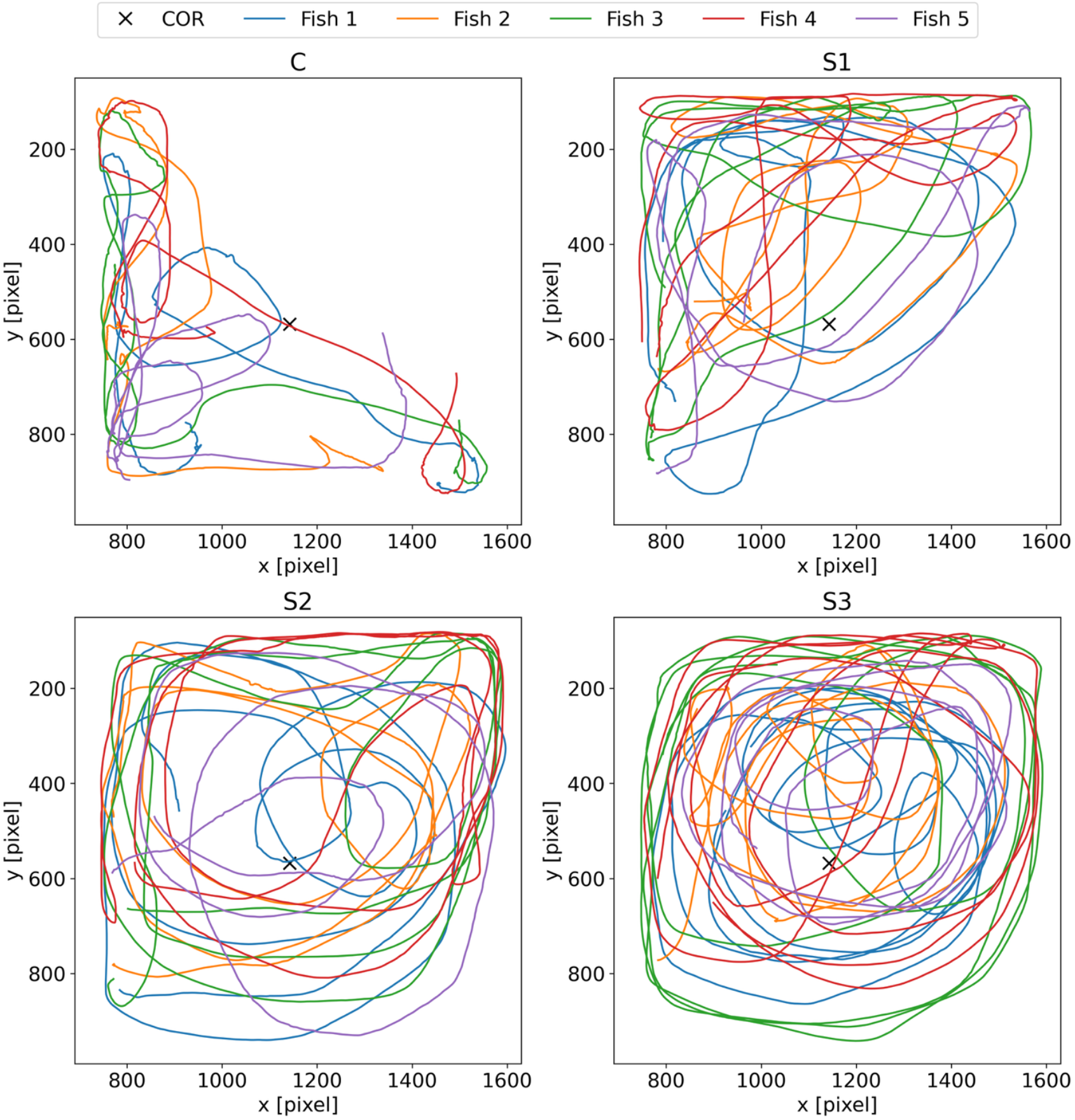
Two-dimensional trajectories of fish across stimulus conditions. Representative trajectories from day 3 (the 2nd cycle) are shown. Under the fastest stimulus condition (S3), fish exhibited rotational movement around the tank centre for most of the period. Control trials showed predominantly exploratory movements near the tank boundaries. Transitional trajectory patterns were often observed under the lower-speed conditions (S1 and S2).

**Supplementary Figure 4:**
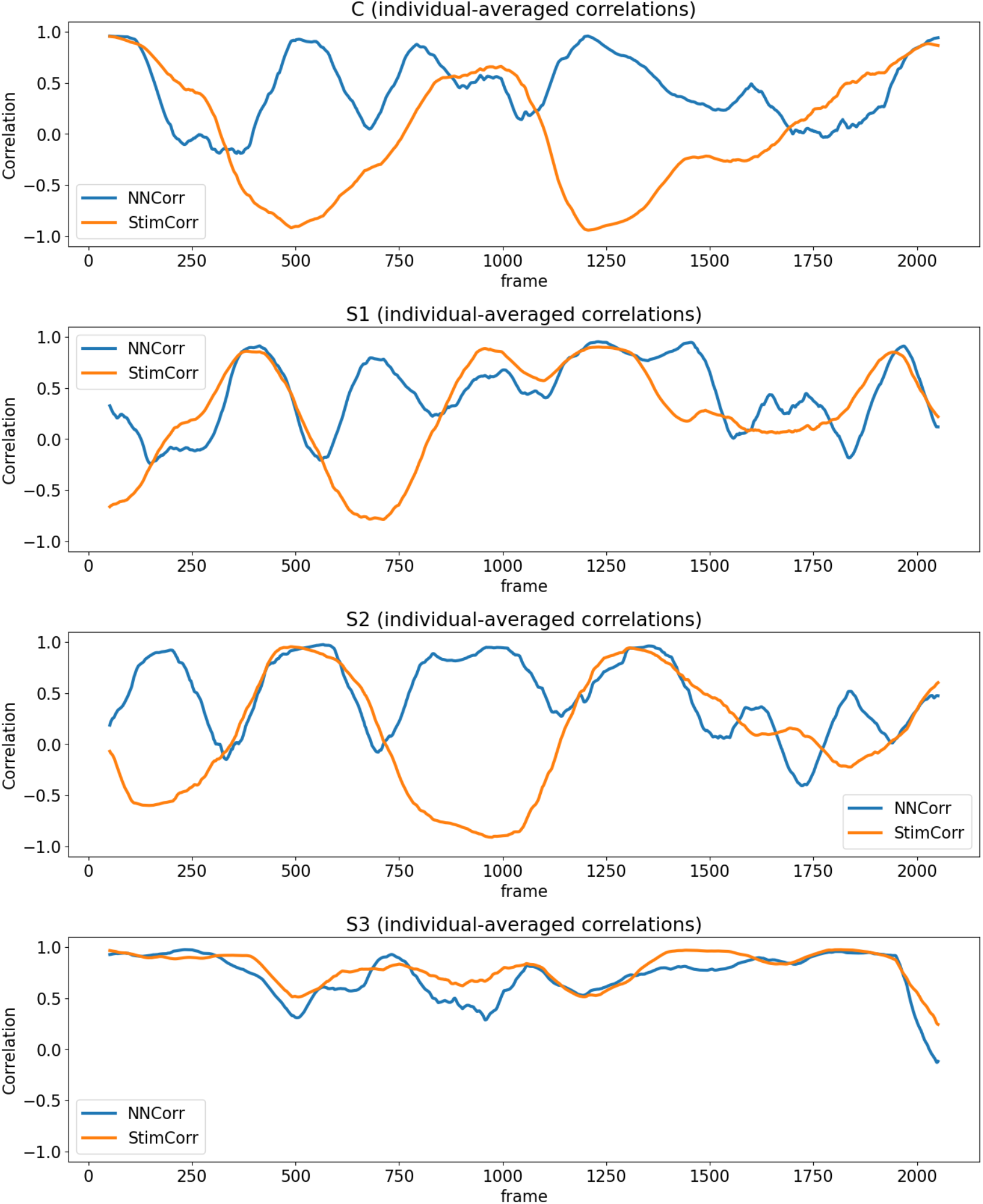
Example of NNCorr and StimCorr dynamics. This example shows the time series of individual-averaged NNCorr (nearest-neighbour directional correlation) and StimCorr (directional correlation with the rotating stimulus) from a representative trial, smoothed using a 100-frame moving average. A similar pattern was observed across other trials.

